# Investigating the Effects of Macaque Primary Motor Cortex Multi-Unit Activity Binning Period on Behavioural Decoding Performance

**DOI:** 10.1101/2020.12.16.423081

**Authors:** Oscar W. Savolainen, Timothy G. Constandinou

## Abstract

This paper investigates the relationship between Multi-Unit Activity (MUA) Binning Period (BP) and Brain-Computer Interface (BCI) decoding performance using Long-Short Term Memory decoders. The motivation is to determine whether lossy compression of MUA via increasing BP has any adverse consequences for BCI Behavioral Decoding Performance (BDP). The Neural data originates from intracortical recordings from Macaque Primary Motor cortex [1]. The BDP is measured by the Pearson correlation *r* between the observed and predicted velocity of the subject’s X-Y hand coordinates in reaching tasks [1]. The results suggest a statistically significant but slight linear relationship between increasing MUA BP and decreasing BDP. For example, when using a 100 ms moving average window, increasing the BP by 10 ms on average reduces the BDP r by approximately 0.85%. This relationship may be due to the reduced number of training examples, or due to the loss of Behavioral information because of reduced MUA temporal resolution.

## I. Background

### A. Wireless Intracortical Brain-Computer Interfaces

Brain Computer Interfaces (BCI) are devices for interfacing electronics with the nervous system. They are generally used to treat neurological conditions. The next generation of Intracortical BCIs is expected to be wireless, due to the infection risks associated with physical transcutanous connections. Due to heating constraints [2], power and thus communication bandwidth are expensive in Wireless Intra-cortical BCIs (WI-BCIs). As such, effective on-implant data compression in WI-BCIs is highly desirable.

### B. Multi-Unit Activity

An intracortical neural signal that is increasing in popularity for BCI behavioral decoding is Multi-Unit Activity (MUA) [3]. When a neuron fires, it creates a sharp voltage spike referred to as an Extracellular Action Potential (EAP) that can be measured by a nearby electrode. MUA signals measure the timing of these EAPs. MUA does not identify the neuron of origin via spike-sorting, as in Single-Unit Activity (SUA). MUA generally outperforms the Local Field Potential (LFP) signal in informativeness [4]. It also outperforms SUA in terms of longevity and ease of computation [4]. However, the effects of the MUA Binning Period (BP) on a BCI decoder’s Behavioral Decoding Performance (BDP) have not yet been determined. This question has significance for the possibility of on-implant data compression [5].

The BP commonly varies from 1-100ms [1], [5]–[7]. The practical lower limit of 1 ms is derived from EAPs lasting approximately 2 ms. Additionally, the firing rate of a neuron typically varies between 10 and 120 Hz [8]. As such, it is unlikely that multiple neurons at an electrode will fire within the same 1 ms window. However, the effects on BDP of increasing BP beyond 1 ms have not yet been determined.

### C. Compressing MUA signals

MUA signals are classically represented by multiplexing channels together. Per time step BP, the length of an MUA signal is classically *n × m,* where *n* is the number of multiplexed channels. The m bits per channel encode in binary the number of events that occurred on that channel in a given BP. *m* is chosen prior to implantation, and generally scales somewhat logarithmically with BP.

Effective lossless compression requires that the observed sequence have a skewed and narrow histogram [9]. This occurs in classically represented MUA signals. It is partly due to the increased likelihood of certain channels having neural events relative to others. Unfortunately, this interchannel probability distribution is not a distribution that can be easily exploited on-implant in WI-BCIs. This is because it would require extensive on-implant memory and calculations, which scale exponentially by a factor of 2^*n×m*^. They are required because prior knowledge of the interchannel distribution before implantation is unavailable.

The other part of the skewed histogram is due to the intra-channel distribution of the number of MUA events per channel per BP. This can be somewhat estimated prior to implantation. There are at least two ways to exploit this distribution that scale with channel count *n*. The first is to give variable length codewords for each symbol, unique to each channel. Therefore each channel uses its own encoder. This requires *n* encoders, and the use of an additional stop signal between each channel to enable full decodability. Each stop symbol has a theoretical minimum length of 1 bit, although in practice this may be larger. Therefore, the minimum condition for compression is the number of stop signals plus the combined intra-channel entropy:

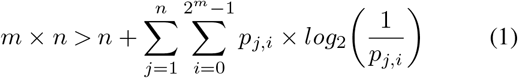

where *p_j,i_* is the probability of *i* events occurring on channel *j*. To take advantage of different intra-channel entropies, each encoder would need to adapt to its channel.

The second method is to use the same encoder for each channel. This is very simple to implement and no interchannel stop signal is required. However, no single channel is guaranteed to have ideal compression. The minimum condition for compression is given by:

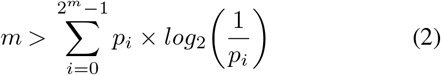

where *p_i_* is the probability of *i* events occurring on the average channel. This only requires a non-uniform distribution for *p_i_*, which is the general, almost-always-met condition for lossless compression. This method is compatible with pre-trained encoders [10], as *p_i_* can be estimated before implantation.

However, if the bandwidth gains from lossless MUA compression are insufficient, MUA signals can also be lossily compressed. By increasing the BP one can achieve lossy compression, at the cost of temporal resolution [5]. Reducing the temporal resolution of MUA potentially affects two things:

- Added BCI latency, which often has a threshold up to which it can be increased without cost, e.g. the update time for a computer screen is approximately 20 ms.
- The decoding performance in BCIs. In this paper, BDP is defined as the Pearson correlation coefficient (r) between the predicted and observed Behavioral data. It is considered a metric of a decoder’s ability to translate the neural information into behavioral output.

### D. Behavioral decoder

The cutting-edge of BCI behavioral decoding involves using Long-Short Term Memory (LSTM) Neural Networks. LSTMs can establish complex non-linear relationships between signals. They also specialize in using information from much earlier on in the sequence to make predictions [11]. The relationship between the Behavioral and Neural data is likely to be non-linear and have an uncertain temporal relationship. Accordingly, LSTMs have been found to be a basis for effective neural decoders [12] [13].

### E. Previous work in evaluating the SUA BP to Behavior relationship

In [13], the authors used various decoders, amongst which LSTMs performed the best, to observe the BDP as a function of SUA BP. The study found that the BDP was independent of the SUA BP between the tested BP range of 1 to 100ms. This suggests that the SUA signal can be significantly compressed without any negative impact on BDP. It would be interesting to determine whether the same BP-to-BDP relationship holds true for MUA.

## II. Methods

This paper investigates the effects of increasing MUA BP on LSTM BDP. A number of different parameters were used to control the BP and other aspects related to the data processing. For all combinations of these parameters, LSTMs were trained and tested on their ability to accurately predict the behavioral information based on MUA input. The other parameters, described further in Section IIB 1, were included to test the effect of BP across various circumstances. All work was done in MATLAB R2019a, using data from [1].

### A. Data description

The dataset is described in [1]. To summarise, the data originated from 12 recording sessions taken across 5 days. Each recording is approximately 12 min long. All data is from Monkey C. The Neural data consisted of SUA recordings from a 96-channel microelectrode array (Blackrock, Inc.) in Macaque Primary Motor cortex. The Behavioral data consisted of the measured hand velocities across the X and Y coordinates, derived from cursor position. The Pearson correlation coefficient (*r*) between the predicted and observed X and Y velocities was used as the BDP metric.

The recordings came in discrete sub-recordings. In the decoding work in this paper, the Neural and Behavioral sub-recordings were respectively concatenated together. This gave better performance than training the decoder on sets of the discrete sub-recordings during parameter optimisation. Only data labelled as having ‘Good’ quality, where the cursor target was reached, were used. In hindsight, this choice was flawed as ‘Bad’ recordings would have been as informative as to the MUA-Behavior relationship.

### B. Processing data

#### 1) Neural data

The neural data was given as SUA sampled at 30 kHz. This was transformed into MUA by concatenating and chronologically sorting the intra-channel SUA firing times. These were then adjusted to the desired BP by binning the number of spiking events within each channel accordingly. The BP was represented by variable *b ϵ* [1:3:20, 30:10:150] ms. In the analysis, it was treated as a continuous variable.

The recording ID *rec* was also tracked, and treated as a categorical variable.

MUA signals, before being fed into decoders, are generally smoothed using a moving average filter, or sometimes using some more complicated statistical measure [14]. This is so as to get a practical estimate of the instantaneous firing rate. It also gives the decoder more non-zero examples to train on. Moving average i.e. boxcar filters were used here, as they are common throughout the literature, perform well and only require one parameter [14]. The filters were causal to be compatible with real-time use. The causal moving average filter was applied to the MUA either once or twice. The number of times smoothed *t ϵ* [12] was treated as a categorical variable. The window width *w ϵ* [50 100:100:500] ms was treated as a continuous variable.

#### 2) Behavioral data

The Behavioral data was originally sampled at 100 Hz. It was resampled using MATLAB’s `resample` function to have a sampling period equal to *b*. Where appropriate, `resample` uses an antialiasing FIR lowpass filter on the input data and compensates for the filter delay. The default value of linear interpolation was used. No other processing was performed on the Behavioral data.

### C. Decoder structure, training and testing

The data was split using a 80-20% training-testing split. The LSTM decoder consisted, in order, of a Sequence Input layer, an LSTM layer with 3 hidden units, a Dropout Layer with 0.2 dropout rate, a Fully Connected layer and a Regression layer. The number of hidden units and dropout rate were optimised on 10% of the training data, i.e. the validation data. The Adam optimization algorithm was used, along with an initial learn rate of 0.1. A learn drop rate of half the number of epochs, and learn drop rate factor of 0.5 were used, based on observation of the training process during validation. The gradient threshold was set to 1.

Each decoder was trained for 150 epochs. This corresponded to a stable but not maximised BDP value, suitable for comparing the effects of the parameters. Computing time was a limiting factor, and this increases approximately linearly with the number of epochs. Decoding performance generally increases logarithmically as a function of epochs.

Each parameter combination was run multiple times to account for any LSTM training stochasticity. The number of successful combinations varied, as some LSTMs failed to train due to time-out or random computation errors. The total number of successful runs was 10071.

After the decoder was trained, it was evaluated using the testing data. The combined processing and decoder training and testing is shown in Fig. 1.

**Fig. 1:**
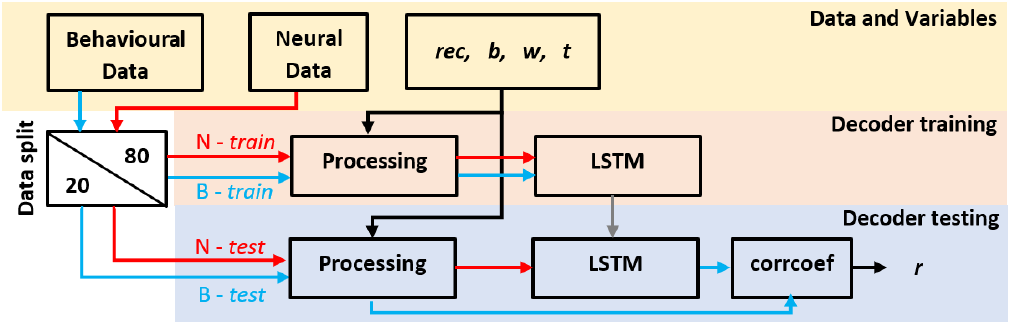
Decoder training and testing diagram.

## III. Results

### A. Combined analysis

The multivariate regression results are given in Table I. This was done using the `fitlm` function. No data was excluded from the analysis. The adjusted R^2^ value was 0.821, *p* = 0. The Durban-Watson test demonstrated significant autocorrelation *(d* = 1.3163, *p* = 1.2e-260). As such, the *p* values observed in Table I are not as significant as shown, if still likely to be extremely significant. Fig. 2 shows a surface plot of the mean BDP as a function of *b* and *w*.

**Fig. 2:**
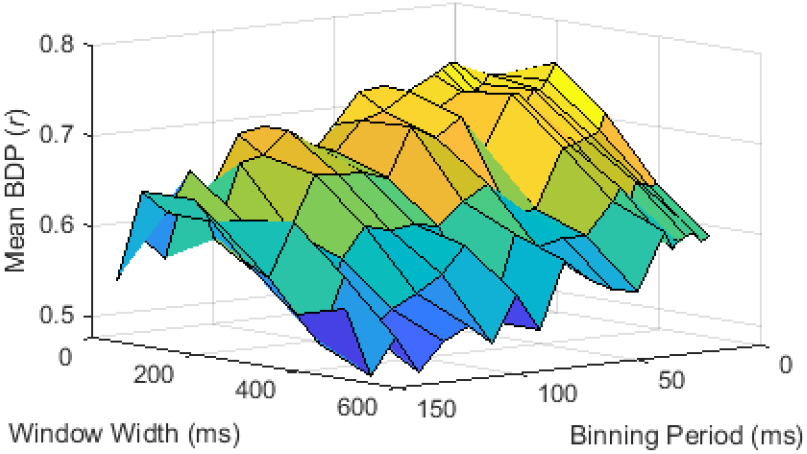
Surface plot of mean Behavioral Decoding Performance as a function of Binning Period *(b)* and Window Width (*w*).

**TABLE I:**
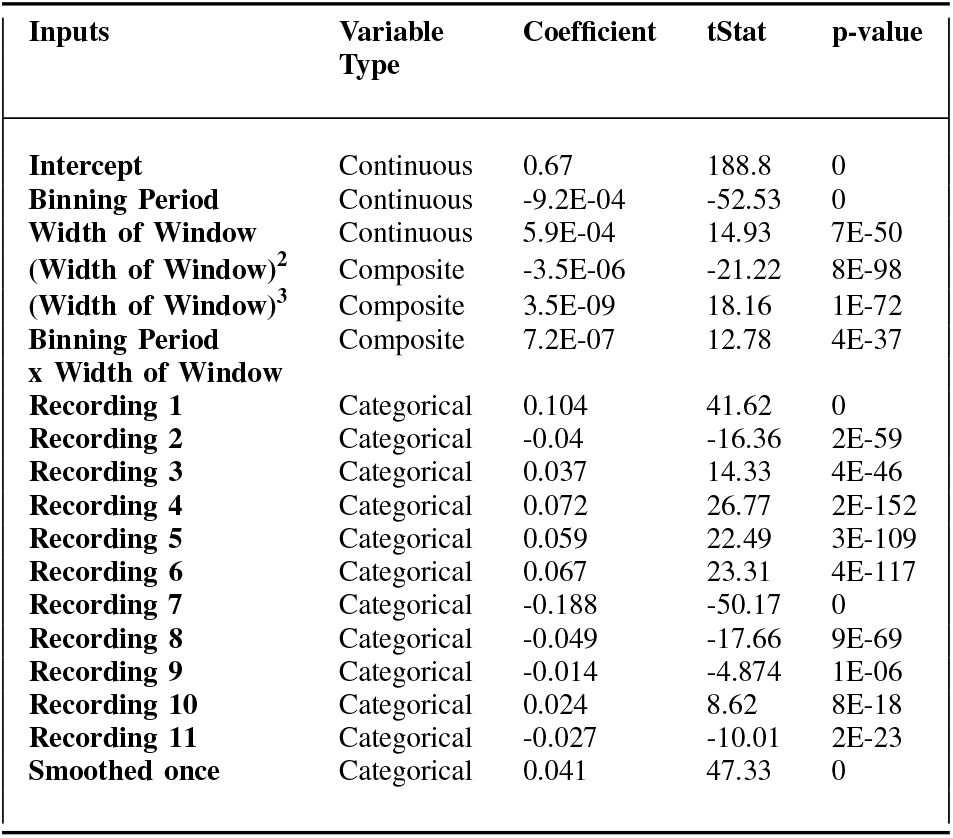
Results from Linear Regression, modeling Behavioral Decoding Performance as a function of Binning Period, Window Width, Recording ID and Times Smoothed.

### B. Binning Resolution

Fig. 3 shows the BDP solely as a function of *b*. The key finding of this paper is that a negative correlation between b and the BDP can be observed in Fig. 2 and 3, and Table I. Another finding is that the mean BDP oscillated around the 1^st^ order linear fit in Fig. 3. This oscillation suggests autocorrelation and the existence of an endogenous term.

**Fig. 3:**
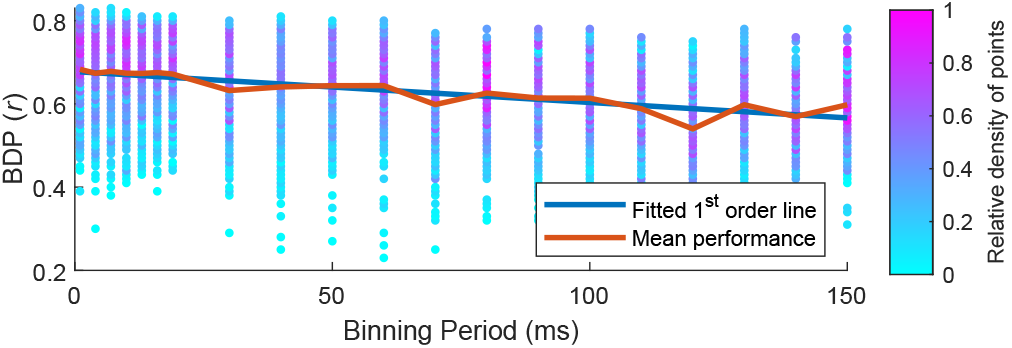
Scatter plot of Behavioral Decoding Performance as a function of Binning Period (*b*). The density of points is colour-coded so as to give some intuition as to the mean weightings.

However, the analysis was unable to find a fit that reduced the autocorrelation to anything below extreme statistical significance. A Fourier Transform of the mean BDP as a function of BP showed peaks at ~ 20 and ~ 40 Hz. However, the Fourier Transform is limited to 50 Hz due to the resolution of the tested BP range. Therefore aliasing is possible. If this oscillation does have a specific frequency, it may be that certain periodic sets of BPs lead to the LSTM being able decode behavior more accurately. This may suggest some common periodicity between MUA and hand movement velocity.

### C. Window Width

Fig. 4 shows the BDP solely as a function of *w*. It seems that, particularly above a *w* of 200 ms, the BDP decreased with increased *w*. Behavioral information encoded in the MUA may have been lost because of reduced temporal resolution. Based on the spread of values and analysis of the residuals, a cubic fit was found to fit the data best.

**Fig. 4:**
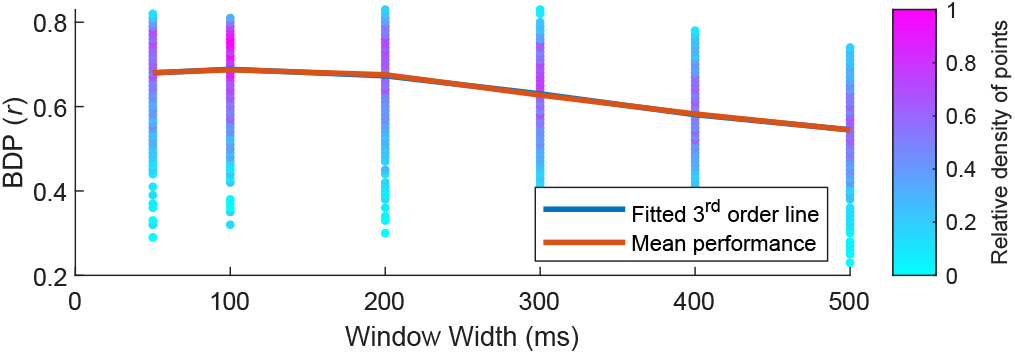
Scatter plot of Behavioral Decoding Performance as a function of Window Width (*w*). The density of points is colour-coded so as to give some intuition as to the mean weightings.

## IV. Discussion

### A. Reduced BDP as a function of BP

The linear decrease in decoding performance as a function of BP may be due to a reduced number of training samples. This is because the number of training samples reduces linearly as a function of BP. To solve this, the number of training samples could be fixed at a certain value for all values of BP. This would require the decoders have sufficient training examples so as to make additional information largely redundant. For example, 10^5^ samples would require a training recording length of approximately 4 hours for a BP of 150 ms, and 2 minutes for a BP of 1ms. With a maximum recording length of 766 s (minimum of 698 s) in this dataset, the lengths may be insufficient to tease out the effects of the number of training samples. Finding the number of required samples would require some parameter optimisation on a large dataset. This warrants some further investigation. However, in cases where large training data sizes are unavailable, we can perhaps practically ignore the mechanism behind the reduced BDP as a function of BP.

As such, in cases with limited training data size, there are potentially two general findings of interest concerning the relationship between BP and BDP. Firstly, the results show an approximately linear relationship between increasing BP and decreasing BDP. For example, from Table I, using a 100 ms window, increasing the MUA BP by 100 ms on average decreases the BDP by 8.5% *r.* This may be an affordable trade-off in cases where the uncompressed data rate is prohibitively large. Alternatively, it may encourage researchers to use smaller bins in cases where data rate is not the limiting factor. Secondly, the autocorrelation, while suggestive of some ideal subset of BP values, is nonetheless minor in amplitude. The oscillation only has a standard deviation of 1.72% *r.* Therefore, it is perhaps something that can be practically ignored when selected an appropriate BP.

### B. Autocorrelation

No other processing was performed other than that specified and controlled for by the input variables. Therefore it seems that the proposed fit between the existing variables and the result is flawed, rather than there existing unaccounted for variables. Additionally, the observed *p* values in Table I are so extreme as to practically guarantee the existence of a relationship when using LSTM decoders. As such, a perfectly statistically characterised relationship between input and output is perhaps not necessary here. The result of interest is more or less clear despite the non-perfect fit.

## V. Conclusion

It is observed that increasing the Binning Period of MUA has a statistically significant negative effect on an LSTM’s Behavioral Decoding Performance. This could be either due to the reduction in number of training examples, or because of the loss of behavioral information, or both. This relationship is not perfectly linear, as suggested by the presence of autocorrelation in the 1^st^ order residual. On average, using a 100 ms moving average filter, increasing the MUA BP by 10ms reduced the BDP by approximately 0.85% *r*.

The results also show that the width of a moving average filter, used on the MUA, generally does not affect the BDP below widths of 200 ms. However, above window widths of 200 ms, increasing the width has a negative effect. This may be because of to the loss of Behavioral information encoded in the MUA data due to reduced temporal resolution.

## VI. Acknowledgements

We thank Robert Flint and Marc Slutzky for making their data available [1].

## References

[1] R. D. Flint et al., “Accurate decoding of reaching movements from field potentials in the absence of spikes,” Journal of neural engineering, vol. 9, no. 4, p. 046006, 2012.

[2] P. D. Wolf and W. Reichert, “Thermal considerations for the design of an implanted cortical brain–machine interface (BMI),” Indwelling Neural Implants: Strategies for Contending with the In Vivo Environment, pp. 33–38, 2008.

[3] D. M. Brandman et al., “Rapid calibration of an intracortical brain– computer interface for people with tetraplegia,” Journal of neural engineering, vol. 15, no. 2, p. 026007, 2018.

[4] E. Stark and M. Abeles, “Predicting movement from multiunit activity,” Journal of Neuroscience, vol. 27, no. 31, pp. 8387–8394, 2007.

[5] N. Even-Chen et al., “Power-saving design opportunities for wireless intracortical brain–computer interfaces,” Nature Biomedical Engineering, pp. 1–13, 2020.

[6] R. Harrison et al., “A low-power integrated circuit for a wireless 100-electrode neural recording system,” in 2006 IEEE International Solid State Circuits Conference-Digest of Technical Papers. IEEE, 2006, pp. 2258–2267.

[7] B. Jarosiewicz et al., “Virtual typing by people with tetraplegia using a self-calibrating intracortical brain-computer interface,” Science translational medicine, vol. 7, no. 313, pp. 313ra179–313ra179, 2015.

[8] B. Gosselin, “Recent advances in neural recording microsystems,” Sensors, vol. 11, no. 5, pp. 4572–4597, 2011.

[9] C. E. Shannon, “A mathematical theory of communication,” Bell Syst. Tech. J., vol. 27, no. 3, pp. 379–423, 1948.

[10] O. W. Savolainen and T. G. Constandinou, “Lossless compression of intracortical extracellular neural recordings using non-adaptive huffman encoding,” in 2020 42nd Annual International Conference of the IEEE Engineering in Medicine & Biology Society (EMBC). IEEE, 2020, pp. 4318–4321.

[11] C. Olah, “Understanding LSTM networks,” 2015.

[12] T. Hosman et al., “BCI decoder performance comparison of an LSTM recurrent neural network and a kalman filter in retrospective simulation,” in 2019 9th International IEEE/EMBS Conference on Neural Engineering (NER). IEEE, 2019, pp. 1066–1071.

[13] J. I. Glaser et al., “Machine learning for neural decoding,” arXiv preprint arXiv:1708.00909, 2017.

[14] S. Koyama et al., “Comparison of brain–computer interface decoding algorithms in open-loop and closed-loop control,” Journal of computational neuroscience, vol. 29, no. 1-2, pp. 73–87, 2010.

